# RangeShifter 2.0: An extended and enhanced platform for modelling spatial eco-evolutionary dynamics and species’ responses to environmental changes

**DOI:** 10.1101/2020.11.26.400119

**Authors:** Greta Bocedi, Stephen C. F. Palmer, Anne-Kathleen Malchow, Damaris Zurell, Kevin Watts, Justin M. J. Travis

## Abstract

1. Process-based models are becoming increasingly used tools for understanding how species are likely to respond to environmental changes and to potential management options. RangeShifter is one such modelling platform, which has been used to address a range of questions including identifying effective reintroduction strategies, understanding patterns of range expansion and assessing population viability of species across complex landscapes.
2. Here we introduce a new version, RangeShifter 2.0, which incorporates important new functionality. It is now possible to simulate dynamics over user-specified, temporally changing landscapes. Additionally, the genetic and evolutionary capabilities have been strengthened, notably by introducing an explicit genetic modelling architecture, which allows for simulation of neutral and adaptive genetic processes. Furthermore, emigration, transfer and settlement rules can now all evolve, allowing for sophisticated simulation of the evolution of dispersal. We illustrate the potential application of RangeShifter 2.0’s new functionality by two examples. The first illustrates the range expansion of a virtual species across a dynamically changing UK landscape. The second demonstrates how the software can be used to explore the concept of evolving connectivity in response to land-use modification, by examining how movement rules come under selection over landscapes of different structure and composition.
3. RangeShifter 2.0 is built using object-oriented C++ providing computationally efficient simulation of complex individual-based, eco-evolutionary models. The code has been redeveloped to enable use across operating systems, including on high performance computing clusters, and the Windows GUI has been enhanced. Furthermore, the recoding of the package has supported the development of a new version running under the R platform, RangeShiftR.
4. RangeShifter 2.0 will facilitate the development of in-silico assessments of how species will respond to environmental changes and to potential management options for conserving or controlling them. By making the code available open source, we hope to inspire further collaborations and extensions by the ecological community.

## Introduction

Faced with an accelerating global biodiversity crisis caused by multiple interacting and often anthropogenic environmental changes (Ceballos et al., 2015; Urban, 2015; IPBES, 2019), biologists are striving to understand and predict how species will respond, in both ecological and evolutionary terms, to these threats and to management interventions (Urban et al., 2016; Urban, 2019). Policy makers, conservation biologists and land managers are relying more and more on such predictions to manage biodiversity on multiple fronts, including protecting threatened species, limiting invasive species, and targeting habitat restoration efforts to both enhance in-situ conservation and promoting range shifting (IPBES, 2019). Process-based models, also called dynamic or mechanistic models, have become increasingly popular following many calls urging the ecological community to move beyond correlative approaches towards models that explicitly incorporate the key processes underpinning eco-evolutionary responses to environmental changes (Franklin, 2010; Huntley et al., 2010; Schurr et al., 2012; Evans et al., 2013; Thuiller et al., 2013; Urban et al., 2016; Cabral et al., 2017; Connolly et al. 2017; Briscoe et al., 2019; Peterson et al., 2019). Several models and platforms are actively being developed (e.g. Lurgi et al. 2015; Landguth et al. 2017; Okamoto & Amarasekare, 2018; Schumaker & Brookes, 2018; Cotto et al. 2020; Kearney & Porter, 2020; Visintin et al., 2020), benefits and shortcomings scrutinised (Dormann et al., 2012; Singer et al., 2016; Zurell et al., 2016; Fordham et al., 2018; Johnston et al., 2019), and a promising variety of applications is emerging (e.g. Synes et al., 2016).

RangeShifter is a process-based models that we initially developed (Bocedi, Palmer, et al., 2014), in response to the many calls for moving towards integrated dynamic modelling approaches. The main objective was to provide an individual-based, spatially-explicit modelling platform that integrated population dynamics with sophisticated dispersal behaviour, and that could be used for a variety of applications, from theory development to in-silico testing of management interventions. Indeed, since its release, RangeShifter has been used in studies addressing a range of issues, including testing the effectiveness of alternative management interventions to improve connectivity and population persistence (Aben et al., 2016; Henry et al., 2017), facilitating range expansion (Synes et al., 2015, 2020), improving reintroduction success (Heikkinen et al., 2015; Ovenden et al., 2019), investigating range dynamics of invasive (Fraser et al., 2015; Dominguez Almela et al., 2020) and recovering species (Sun et al., 2016) and theoretically investigating how different traits and processes affect rate of range expansion (Bocedi, Zurell et al. 2014; Henry et al., 2014; Barros et al., 2016; Santini et al., 2016). RangeShifter has also been coupled with CRAFTY (Murray-Rust et al., 2014), an agent-based model designed to explore the impact of land managers’ behaviours on land-use change, showing that, in the example context of predicting interactions between crops and their pollinators in a changing agricultural landscape, models that integrate ecological processes with land managers’ behaviours, together with their interactions and feed-backs can reveal important dynamics in land use change which might otherwise be missed (Synes et al., 2019; Willemen et al., 2019).

Here, we present the new RangeShifter 2.0, which, among various additions and improvements, includes two major novelties: the option for implementing temporally dynamic landscapes and a module for the explicit modelling of neutral and adaptive genetics (controlling dispersal traits). RangeShifter is written in C++; it has been completely recoded from its original release following object-oriented programming principles and is now open source, thus facilitating wider usage and enhancements by the ecological community. Additionally, we provide a dedicated website (https://rangeshifter.github.io/) and updated tutorials for learning to use RangeShifter, and a forum page for more effective communication among users. In addition to the original and improved Windows graphical user interface (GUI), RangeShifter can now be compiled to run in batch-mode on Linux computer clusters. Below we briefly describe, and illustrate with examples, the two major additions of dynamic landscapes and explicit genetics, while we refer to the RangeShifter 2.0 User Manual (https://github.com/RangeShifter/RangeShifter-software-and-documentation) for smaller changes and new features.

## Model enhancements

### DYNAMIC LANDSCAPES

Considering dynamically changing landscapes is crucial for scenario-based simulations (e.g. climate change or land-used change scenarios), for implementing landscape processes through time (e.g. ongoing habitat fragmentation) and for testing dynamic management interventions accounting for time lags from their deployment (e.g. creating a new woodland) to the realization of their full potential (Watts et al., 2020). In RangeShifter 2.0, the landscape may be changed any number of times during a simulation, but always at the start of the year, i.e. prior to reproduction. The changes may comprise any of: alterations to the habitat structure; addition, removal or changes of patches in a patch-based model; and modifications of the cost map when using the stochastic movement simulator (SMS; Palmer, Coulon, & Travis, 2011).

### EXPLICIT GENETICS

A new module is provided to define the genetic architecture of a species in a flexible and explicit way. Individuals may carry one or more chromosomes, to which neutral loci and adaptive loci controlling dispersal traits are mapped. It is possible to model unlimited neutral markers, thus allowing tracking of population structure and neutral diversity, as well as simulating spatial genetic patterns emerging from the interaction between demographic and spatial processes, e.g. for in-silico applications of landscape genetics (Manel et al., 2003). The dispersal traits have been extended to cover density-dependent emigration and settlement reaction norms, which may optionally differ between the sexes. Additionally, if SMS is selected as the movement model in the transfer phase, the parameters controlling directional persistence and the dispersal bias and its decay (see the User Manual) can be modelled as evolving traits. Each dispersal or movement trait can be controlled by a separate single chromosome, akin to RangeShifter v1 (Bocedi, Palmer, et al., 2014), or through a highly flexible mapping of traits to chromosomes, which enables the degree of linkage between traits to be controlled and, optionally, pleiotropy to be incorporated, thus allowing for complex genetic architectures underlying evolution of dispersal strategies (Saastamoinen et al., 2018). The whole genome of each individual may be output in a separate file if required, e.g. for the calculation of landscape genetic indices.

## Example applications

### EFFECTIVENESS OF WOODLAND CREATION STRATEGIES TO FACILITATE RANGE EXPANSION

We illustrate the application of dynamic landscapes using the example of woodland creation in a real UK landscape introduced by Synes et al. (2015, 2020), who compared the effects of various realistic management scenarios for improving functional connectivity for a range of exemplar virtual woodland species on both species’ persistence in existing patches and range expansion ability. They compared persistence and expansion rates under the management scenarios with a baseline rate for the original landscape. However, as the landscape changes were ‘instant’, i.e. the new habitat was assumed to be immediately fully suitable, the differences they observed could be over-estimated, as newly planted woodland would in reality take many decades to develop into the equivalent of existing woodland in terms of its suitability as breeding habitat for many species (Watts et al., 2020). Rather, newly planted areas might be expected firstly to provide increased structure which might aid movement of woodland species, and then gradually increase in quality as breeding habitat as canopy cover develops.

We assume as in Synes et al. (2015, 2020) that the locations of all new woodlands are allocated immediately and on land previously used as improved grassland or arable, and that planting of saplings occurs instantly in all locations. However, rather than instantly becoming mature woodland habitat, planted areas develop gradually over a period of 40 years (Table 1, Fig. 1). We compared the dynamic landscape approach with the ‘instant’ landscape approach on the basis of the two most successful scenarios identified by Synes et al. (2015), namely ‘CreateRandom’ (new patches created anywhere) and ‘CreateSmallAdjacent’ (new planting to increase the size of existing patches of under 3 ha), applied to 4% of the landscape. For illustrative purposes we consider one virtual woodland species with simple sexual, stage-structured demography and good dispersal abilities (Bird_D^+^P^-^S^+^ in Synes et al. 2015; see Table S1 for the model parameters). We modelled dispersal movements through the landscape with SMS. To ensure that the species was in equilibrium before management commenced, we ran simulations for 50 years on the original landscape before applying the first landscape change, and then continued for a further 100 years during which range expansion was allowed to occur under the management scenario. Simulations were run on the baseline landscape and on all the 10 replicate landscapes for each scenario generated by Synes et al. (2015), and each simulation was replicated 10 times.

**Figure 1.**
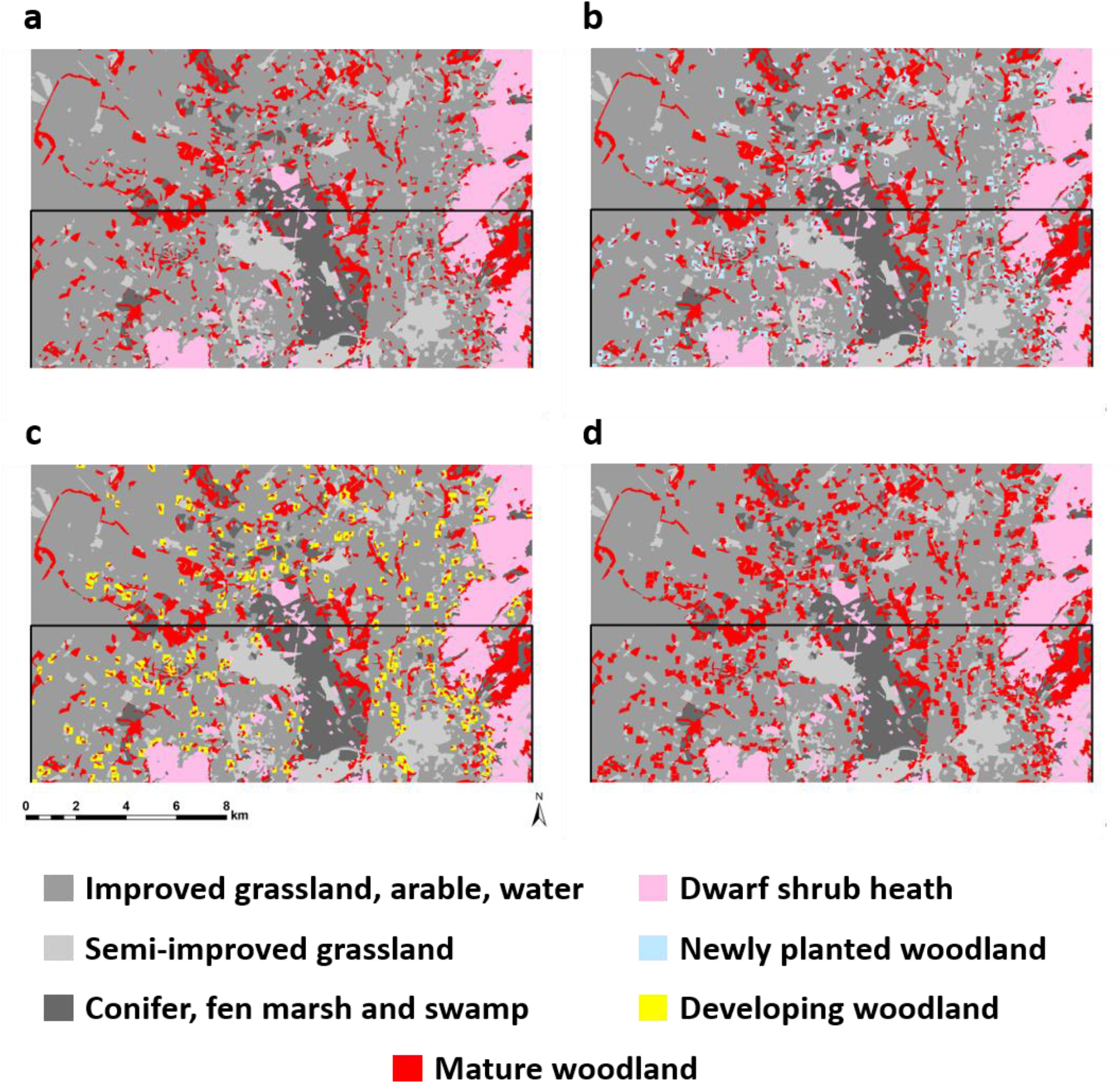
Example of dynamic landscape development: (a) initial landscape, (b) after 5 years when newly planted woodland adjacent to small patches is treated as dwarf shrub heath for dispersal modelling, (c) after 20 years as canopy closure develops, (d) final landscape after 40 years when newly planted woodland is fully mature. The black line shows the northern limit of the initial range.

**Table 1.**
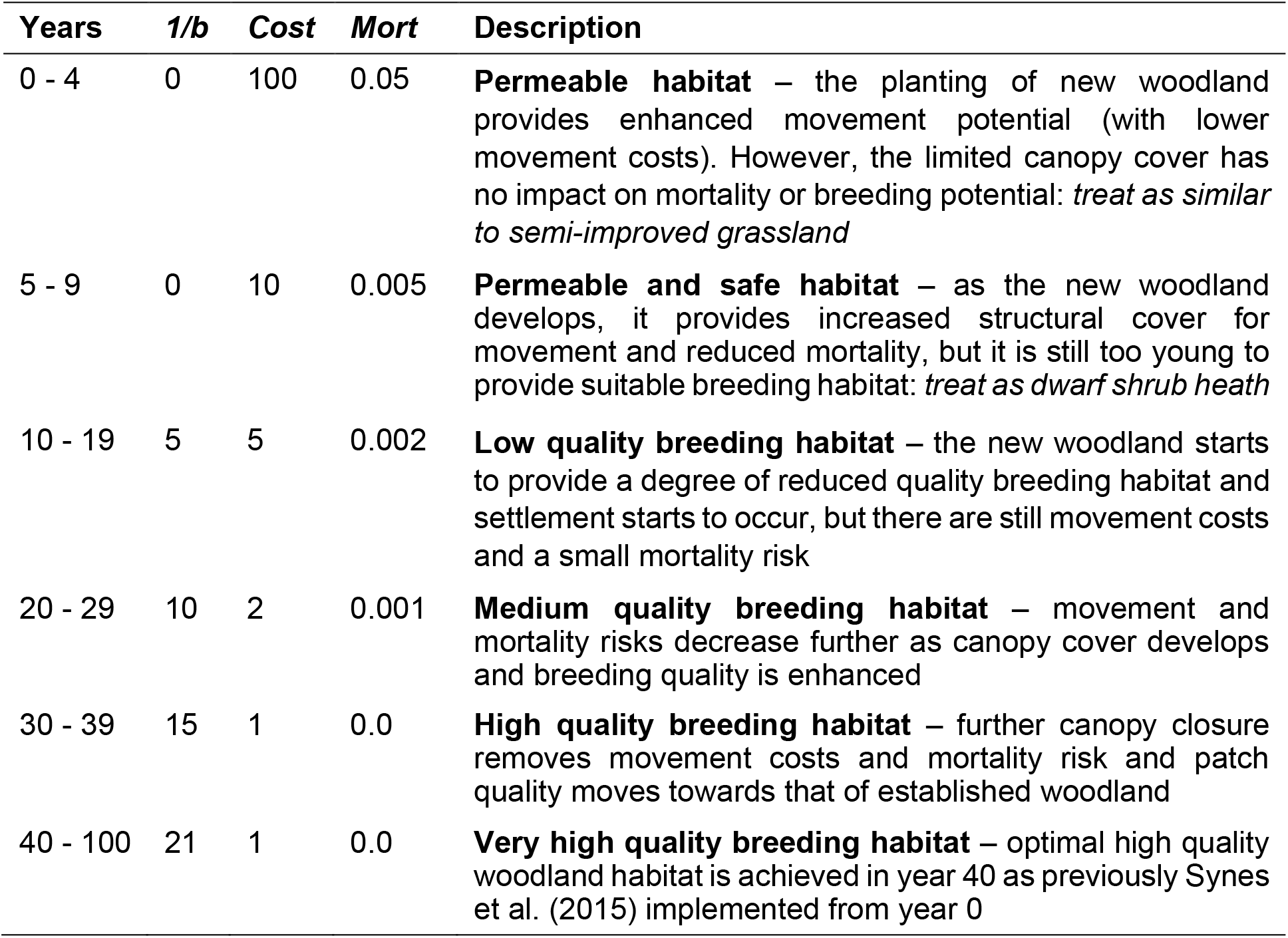
Temporal development of newly planted woodland, where *1/b* is the fecundity density-dependent coefficient (individuals / ha) (which largely determines the equilibrium density of the population), *Cost* is the perceived movement cost applied in modelling the transfer phase of dispersal by SMS and *Mort* is the per-step habitat-specific mortality probability.

For the instant landscape approach (Synes et al. 2015), the mean rate of range expansion for the CreateRandom scenario was 187 m/year over 100 years (standard error s.e. 2.34 m/year), 2.0 times the rate on the baseline landscape. Similarly, for the CreateSmallAdjacent scenario, the mean rate of range expansion was 201 m/year (s.e. 2.90 m/year), 2.1 times faster than the baseline. By applying the dynamic landscape approach to the CreateRandom scenario, the mean rate of range expansion was reduced negligibly to 184 m/year (s.e. 2.64 m/year; relative reduction 1.6%). In contrast, for the CreateSmallAdjacent scenario, the mean rate of range expansion was increased slightly to 216 m/year (s.e. 2.69 m/year; relative increase 7.5%). Despite rather similar total expansion rates over a period of 100 years, the temporal trajectories differed considerably between the instant and the dynamic landscape approach, as is illustrated for a single landscape replicate of the CreateSmallAdjacent scenario (Fig. 2). The total population size on the dynamic landscape lagged behind that on the instant landscape by up to 25% during the first 40 years after planting (Fig. 2A), and the location of the northern range margin on the dynamic landscape was up to 5 km further south during the succeeding 40 years (Fig. 2B).

**Figure 2.**
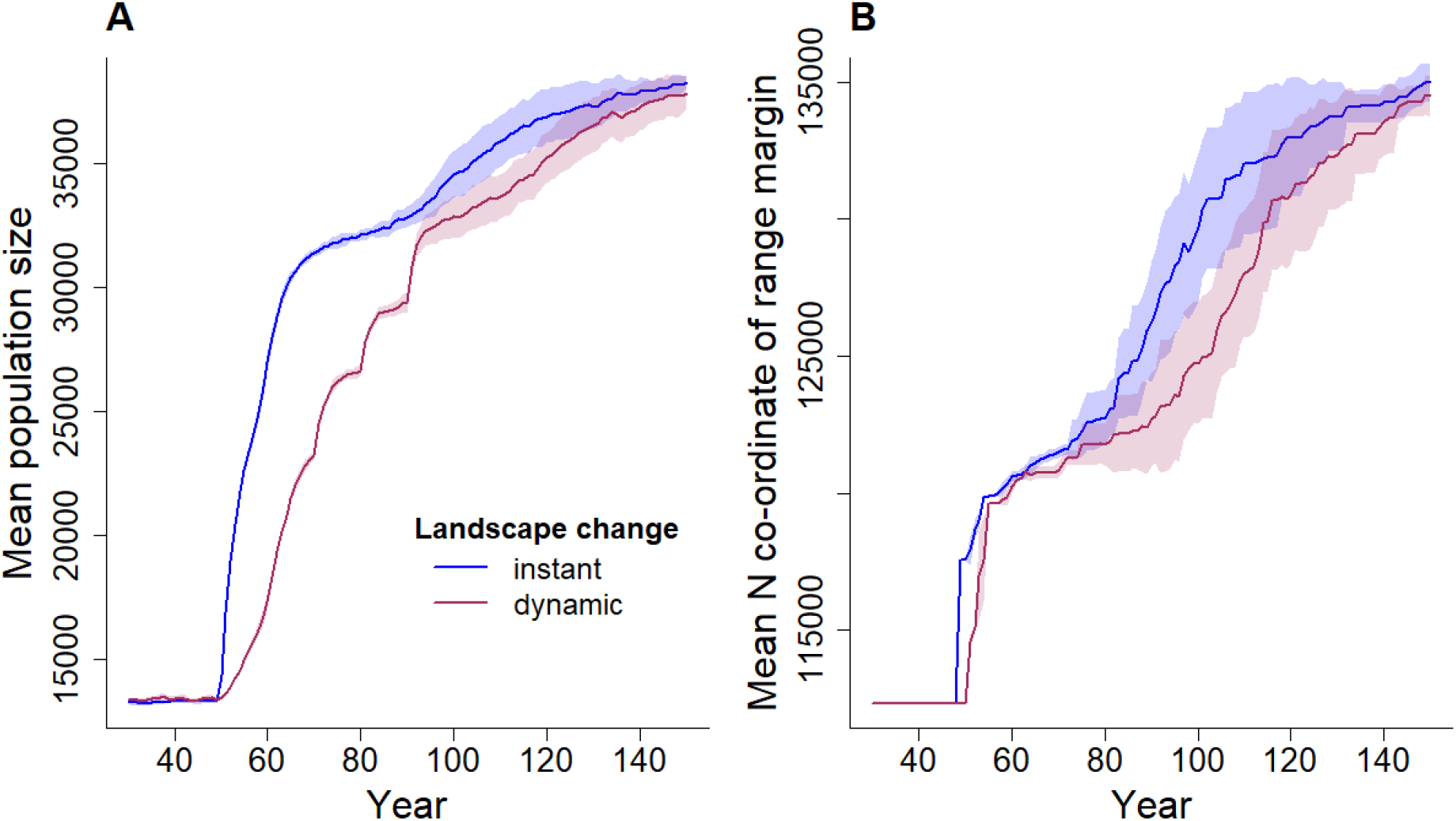
Consideration of dynamic landscape restoration affects predictions on species’ range expansion dynamics. (A) Mean total population size and (B) mean location of species’ northern range margin for the instant (blue) and dynamic (red) landscape change methods commencing at year 50 for a single landscape replicate of the CreateSmallAdjacent scenario. Shades show 95% confidence intervals from 10 replicates.

### EVOLUTION OF MULTIPLE DISPERSAL TRAITS

We illustrate how RangeShifter 2.0 can be used to model evolution of complex dispersal strategies, which involve evolution of multiple traits defining all three phases of dispersal (emigration, transfer through the landscape and settlement in a new habitat patch) on landscapes that differ in their structure and composition. We modelled the evolution of dispersal traits of an annual sexual species on a set of three stylised landscapes of 121 rows x 121 columns differing in the degree to which movement was inhibited by the presence of high-cost cells in the landscape (Fig. 3). Temporally uncorrelated local environmental stochasticity was applied in two forms in order to promote dispersal evolution: as annual variability in carrying capacity and as a small probability of local patch extinction. The parameters controlling all three phases of dispersal, namely emigration (density-dependent), transfer and settlement (density-dependent), evolved independently of one another, each trait being determined by a separate autosome having three unlinked loci. Emigration and settlement traits were modelled as sex-dependent, thus having sex-limited phenotypic expression. Further, sex differences were assumed in settlement: while for both sexes settlement probability could evolve density dependence, males had the additional fixed settlement condition of requiring the presence of a mate in the patch.

**Figure 3.**
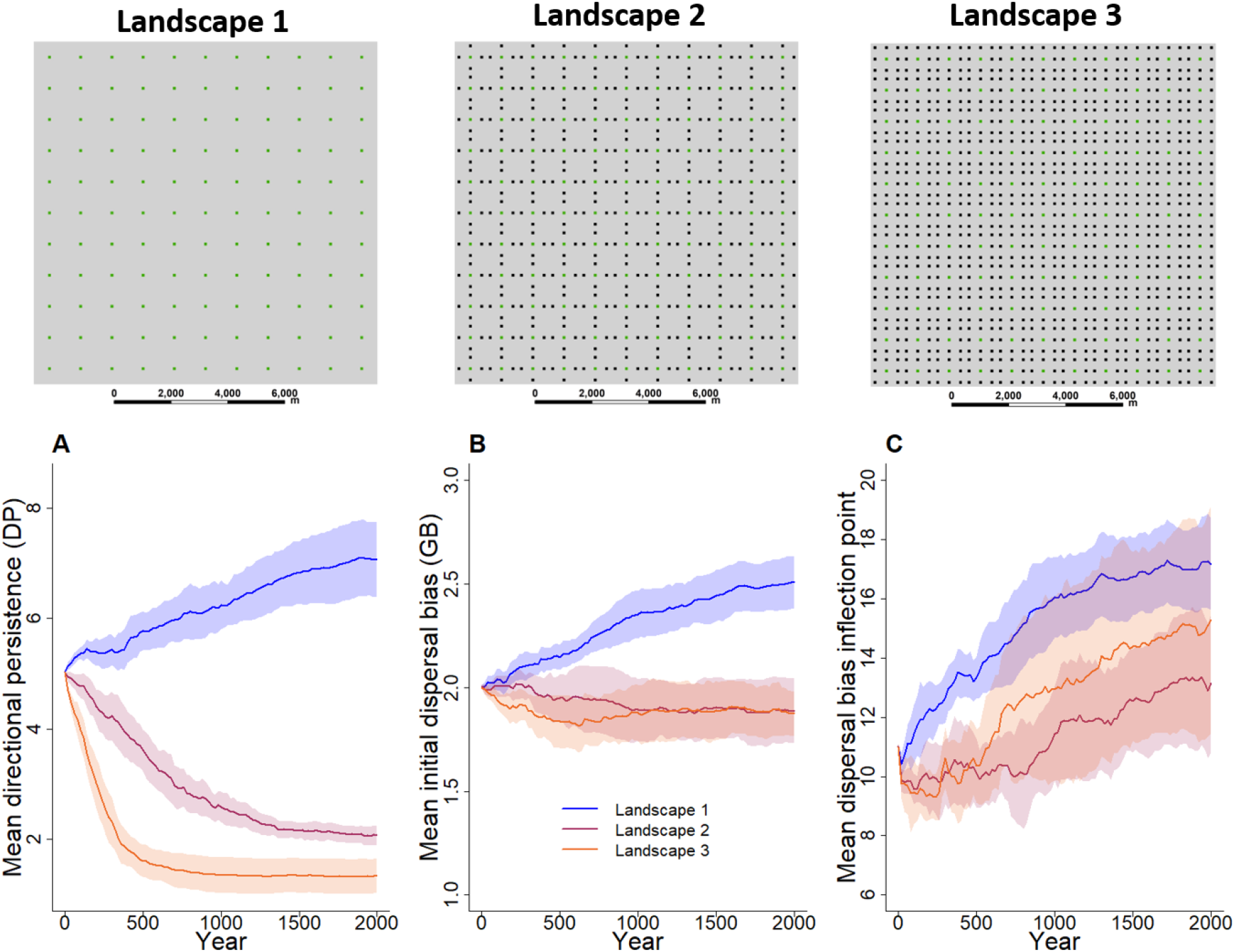
Stylised landscapes used to model evolution of dispersal traits (upper panels). Landscape 1 comprises evenly distributed breeding habitat patches of 100 m x 100 m (green) set in a homogenous matrix (grey). In Landscape 2 high-cost cells inhibitory to movement (black) are added orthogonally between the patches. In Landscape 3 additional inhibitory cells are added to the diagonals between patches. **(A-C)** Evolution of mean transfer traits, directional persistence (A), initial dispersal bias (B) and dispersal bias inflection point (measured in steps taken; C), on the three landscapes. Phenotypic values are averaged over all individuals and 10 replicate simulations.

The transfer phase of dispersal was modelled using SMS, which models the dispersal trajectory on a stepwise basis whilst accounting for perceived movement costs and a tendency to follow a correlated path, as determined by the directional persistence (DP) and the dispersal bias parameters. The dispersal bias determines the tendency of moving in a straight line away from the natal patch and is subject to a decay in strength as a function of the accumulated number of steps taken (see Supplementary Information). DP and the parameters defining the decay function of the dispersal bias were modelled as evolving traits, thus allowing for evolution of movement rules. In the baseline Landscape 1 (Fig. 3), there were no inhibitory cells in the matrix (Cost = 10; per-step mortality = 0.01), and therefore we would expect relatively straight movement to evolve (Zollner & Lima, 1999). However, in Landscape 2, it is much less clear what would be the best movement strategy, as the orthogonal paths between patches are inhibited by high-cost cells (Cost = 1000; per-step mortality = 0.5), whereas the diagonal movements are not. Finally, in Landscape 3, both orthogonal and diagonal paths are impeded, and dispersers must therefore evolve strategies to reach a new patch whilst avoiding as much as possible the high-risk inhibitory cells. We ran ten replicate simulations of 2000 years on each landscape. All model parameters and initialisation conditions are reported in Table S2. Equations defining reaction norms are also reported in the Supplementary Information.

As expected from the spatial and temporal configuration of the selective environments, the dispersal strategies that evolved on the different landscapes differed mainly in their movement rules (Fig. 3A-C), whereas they evolved similar reaction norms for the emigration and settlement phases (Figs. S1-S2, S4-S5). In the absence of inhibitory features in Landscape 1, very straight movement trajectories evolved (Fig. S6A): both mean directional persistence (DP) and mean dispersal bias (the tendency to move in a direction away from the natal patch) reached high values of ~ 7.0 and 2.5 respectively after 2000 years (Fig. 3A-B), and indeed there was some indication that they were still increasing slightly. In contrast, when orthogonal movement became inhibited in Landscape 2, much less direct movement evolved (Fig. S6B), as determined by low mean DP (Fig. 3A). Mean dispersal bias initially remained relatively high at around 2.0 (Fig. 3B), but its mean inflection point (the number of steps at which dispersal bias decreases most rapidly) decreased from around 16 steps on Landscape 1 to around 12 steps (Fig. 3C). Thus, dispersers having evolved in Landscape 2 would be following a much less straight path sooner after having left the natal patch compared to dispersers having evolved in the more benign Landscape 1 (Fig. S3), thereby enabling them to respond to the appearance of a suitable (low cost) cell within the perceptual range by moving towards it. The addition of inhibitory features to diagonal movement further developed this trend: dispersal bias altered little, but DP decreased to a very low level of around 1.3 on average (Fig. 3A).

Emigration probability generally evolved to be male-biased. Mean male emigration probability decreased as the occurrence of inhibitory cells in the landscape increased (Landscapes 2 and 3) because the cost of dispersal effectively increased (Fig. S1-S2). Male-biased emigration would be expected, given the loosely polygynous mating system (i.e. males can mate with multiple females but each female mates only with one male) and the high environmental and demographic stochasticity (Table S2), which increase between-patch variance in male reproductive success (Henry et al., 2016; Li & Kokko, 2019). Density-dependent settlement evolved similarly in the two sexes, so that individuals were likely to settle at the first suitable patch encountered unless it was substantially above carrying capacity (Fig. S4-S5).

## Discussion

RangeShifter 2.0 provides enhancements and substantial extensions to the RangeShifter software (Bocedi, Palmer, et al., 2014) expanding its potential range of applications. The flexible and spatially-explicit demography and dispersal modules that are distinctive of this platform (Lurgi et al., 2015) can be now combined with a flexible genetically-explicit representation of neutral markers and/or multiple dispersal traits, allowing for diverse applications focussed on combining population genetic processes with ecological and environmental processes (Manel et al., 2003; Epperson et al., 2010) and accounting for evolution of complex and multi-trait dispersal strategies (Cote et al., 2017; Legrand et al., 2017; Saastamoinen et al., 2018). This is further combined with the ability of incorporating dynamic landscapes, enabling applications that explicitly aim to predict species’ genetic, ecological and evolutionary responses to ongoing environmental changes. Such applications include in-silico testing of management interventions which need to account for the occurrence of ecological time-lags when targeting and evaluating conservation actions (Watts et al., 2020).

Importantly, and in contrast with the previous release, RangeShifter 2.0 source code is now open source (https://github.com/rangeshifter), published under the GNU general public license (GPLv3). It is hence free for the wider community to use, modify and share. Furthermore, RangeShifter 2.0 is also the core of the new package RangeShiftR (Malchow et al., 2020), which allows running RangeShifter from the R environment (R Core Team, 2020) while maintaining the high performance of the C++ code, and includes functions assisting with the set-up of the simulations, the parameterisation and output analyses. RangeShiftR, in addition to improving and broadening RangeShifter accessibility, makes it easily available for multiple platforms, has access to R’s infrastructure for parallel and cluster computing and offers many opportunities for interoperation with other R packages.

RangeShifter 2.0 additionally comes with an enhanced Windows graphical user interface (GUI) as freeware (https://github.com/RangeShifter/RangeShifter-GUI). From current users, and from workshops that we are running worldwide, we are able to appreciate the value of the RangeShifter GUI: it is particularly useful for non-modellers to explore eco-evolutionary dynamics and their conservation implications, to recognise data gaps in empirical systems, to communicate with stakeholders, and for teaching purposes across grades. It also provides easily accessible and free software for countries with little funding for conservation and research. Further, to improve accessibility, the User Manual has now been translated into Spanish (https://github.com/RangeShifter/RangeShifter-software-and-documentation).

RangeShifter is in continuous development, and there are key areas for future progress, which we hope, by making it open source and integrating it with R, will be addressed by a common effort to move towards a fully-integrated dynamic platform that includes all the key and necessary processes for predicting species’ eco-evolutionary responses to global changes. For example, RangeShifter 2.0 currently remains a single-species model, while inter-specific interactions are often key in determining species’ persistence to global changes (Gilman et al., 2010; Norberg et al., 2012; Urban et al., 2012; Urban et al., 2019; Bocedi et al., 2013; Svenning et al., 2014; Thompson & Fronhofer, 2019). Although we made a first important step in including explicit genetics, and we are actively prioritising this area of development, RangeShifter 2.0 does not yet include the level of sophistication that characterises much forward-time population genetic software (Guillaume & Rougemont, 2006; Haller & Messer, 2019), in terms of genetic processes, structures and outputs, and adaptive traits. For example, the possibility of modelling adaptation to multiple environmental variables will be a crucial addition. However, RangeShifter 2.0 holds an advantage in terms of the ecological, demographic and dispersal complexity it can represent, which, combined with explicit genetics, opens possibilities for sophisticated landscape genetics applications and for fully accounting for evolution of dispersal behaviours (not just emigration rates) which are likely to be critical for species’ inhabiting or moving through complex, human-modified landscapes.

## Acknowledgments

Development of RangeShifter 2.0 was supported by the project PROBIS funded by the BiodivERsA European Joint Call 2012-2013. We thank R. L. Allgayer, A. Ponchon and N. W. Synes for their help and contribution to the RangeShifter development and application. We also thank the many users of RangeShifter and participants to workshops for their invaluable feedback. GB was supported by a Royal Society University Research Fellowship. AKM and DZ were supported by Deutsche Forschungsgemeinschaft (DFG) under grant agreement No. ZU 361/1-1.

## Author’s contributions

GB, SCFP and JMJT mainly developed the model structure, implemented the C++ core code, developed the GUI and wrote the documentation. AM contributed to code testing and cleaning, and DZ developed the accompanying website. KW provided the landscape data for the first example. GB, SCFP and JMJT wrote the initial manuscript and all authors contributed critically to the drafts and gave final approval for publication.

## Data Availability

RangeShifter C++ core code for the batch implementation is open source under the GNU general public license (GPLv3). The code, as well as the compiled Windows package, the User Manual and the data for the tutorials are available from GitHub https://github.com/RangeShifter.

