## Supplementary Material for "RangeShifter 2.0: An extended and enhanced platform for modelling spatial eco-evolutionary dynamics and species’ responses to environmental changes"

EFFECTIVENESS OF WOODLAND CREATION STRATEGIES TO FACILITATE RANGE EXPANSION

**Table S1**. RangeShifter parameters and options applied in the dynamic landscape example.

| Parameter / option | Value | | |
|---|---|---|---|
| ***Demographic: stage-structured sexual*** | | | |
| Proportion of males | 0.5 | | |
| No. of stages | 2 | | |
| Maximum age (years) | 5 | | |
| Maximum mean fecundity (young / female) | 5.0 | | |
| Juvenile survival rate | 1.0 | | |
| Adult survival rate | 0.5 | | |
| Habitat-specific density dependence (*1/b*) | See Table 1 | | |
| ***Emigration: stage- and density-dependent*** | ***d0*** | ***alpha*** | ***beta*** |
| Juveniles | 0.5 | 3.0 | 0.5 |
| Adults | 0.0 | 0.0 | 0.0 |
| Initial inflection point (*beta*) females and males | 1.0 | 0.5 | 0.5 |
| ***Transfer: stochastic movement simulator*** | ***cost*** | ***per-step mortality*** | |
| Improved grassland / arable / water | 1000 | 0.05 | |
| Semi-improved grassland | 100 | 0.025 | |
| Conifer, fen marsh and swamp | 25 | 0.005 | |
| Dwarf shrub heath | 10 | 0.001 | |
| Woodland habitats | See Table 1 | | |
| Perceptual range (cells) | 20 | | |
| Perceptual range method | 1 (arithmetic mean) | | |
| Directional persistence (*DP*) | 15.0 | | |
| Memory size (cells) | 1 | | |
| Goal type | 0 (none) | | |
| Straighten path after decision not to settle | yes | | |
| ***Settlement: sex-dependent*** | | | |
| Mate required | only for males | | |
| ***Initialisation: free, all suitable cells at half carrying capacity*** | | | |
| ***Freeze initial range until year 50*** | | | |

EVOLUTION OF MULTIPLE DISPERSAL TRAITS

**Table S2**. RangeShifter parameters and options applied for modelling evolution of multiple dispersal traits.

| Parameter / option | Value | | |
|---|---|---|---|
| ***Landscape*** | | | |
| Environmental stochasticity | in *K* | | |
| Spatial autocorrelation | local | | |
| Temporal autocorrelation | 0.0 | | |
| Amplitude | 0.5 | | |
| Minimum *K* | 10.0 | | |
| Maximum *K* | 50.0 | | |
| Local extinction probability | 0.015 | | |
| ***Demographic: non-structured sexual*** | | | |
| Proportion of males | 0.5 | | |
| Maximum mean fecundity ($R_{max}$) | 10.0 | | |
| Competition coefficient (*bc*) | 1.0 | | |
| Carrying capacity (*K*) | 30.0 | | |
| ***Emigration: sex- and density-dependent*** | | | |
| ***evolving parameters*** | ***mean*** | ***s.d.*** | ***scaling*** |
| Initial maximum prob. (*D0*) females and males | 0.5 | 0.3 | 0.3 |
| Initial slope (*alpha*) females and males | 5.0 | 3.0 | 3.0 |
| Initial inflection point (*beta*) females and males | 1.0 | 0.5 | 0.5 |
| ***Transfer: stochastic movement simulator*** | | | |
| Cost / per-step mortality of suitable breeding | 1 / 0.0 | | |
| Cost / per-step mortality of matrix habitat | 10 / 0.01 | | |
| Cost / per-step mortality of inhibitory habitat | 1000 / 0.5 | | |
| Perceptual range (cells) | 2 | | |
| Perceptual range method | 1 (arithmetic mean) | | |
| Memory size (cells) | 3 | | |
| Goal type | 2 (dispersal bias) | | |
| ***evolving parameters*** | ***mean*** | ***s.d.*** | ***scaling*** |
| Initial directional persistence (*DP*) | 5.0 | 2.0 | 2.0 |
| Initial goal bias (*GB*) | 2.0 | 0.5 | 0.5 |
| Initial dispersal bias slope (*alphaDB*) | 0.5 | 0.25 | 0.25 |
| Initial dispersal inflection point (*betaDB*) | 10 | 10 | 10 |
| ***Settlement: sex- and density-dependent*** | | | |
| Mate required | only for males | | |
| ***evolving parameters*** | ***mean*** | ***s.d.*** | ***scaling*** |
| Initial maximum prob. (*S0*) females and males | 0.5 | 0.25 | 0.25 |
| Initial slope (*alphaS*) females and males | -5.0 | 3.0 | 3.0 |
| Initial inflection point (*betaS*) females and males | 1.0 | 0.5 | 0.5 |
| ***Genetics: one chromosome per trait, 3 loci per chromosome*** | | | |
| Mutation probability | 0.0001 | | |
| Crossover probability | 0.5 | | |
| Initial allele s.d. | 0.1 | | |
| Mutation s.d. | 0.1 | | |
| ***Initialisation: free, all suitable cells at carrying capacity*** | | | |

***Evolution of density-dependent emigration reaction norms***

The emigration probability for individual *i* is given by (see User Manual, section 2.5.1):

$$d_i = \frac{D_{0,\mathrm{i}}}{1 + e^{-\left(\frac{N_{x,y}}{K_{x,y}} - \beta_i\right)\alpha_i}}$$

where $D_{0,i}$, $\beta_i$ and $\alpha_i$ are the individual's phenotypes for the three evolving traits representing: the maximum emigration probability ($D_0$), and the function inflection point ($\beta$) and slope at the inflection point ($\alpha$). $N_{x,y}$ and $K_{x,y}$ represent the patch total number of individuals and carrying capacity respectively.

Emigration generally evolved to be male-biased (Fig. S1). Males evolved a density-dependence emigration strategy by which they start emigrating at lower density (lower inflection point $\beta$, Fig. S2D) than females and have overall higher emigration probabilities than females (higher maximum emigration probability $D_0$, Fig. S2C). As the occurrence of inhibitory cells in the landscape increased (Landscapes 2 and 3), mean male emigration probability decreased (Fig. S1B), as the cost of dispersal effectively increased.

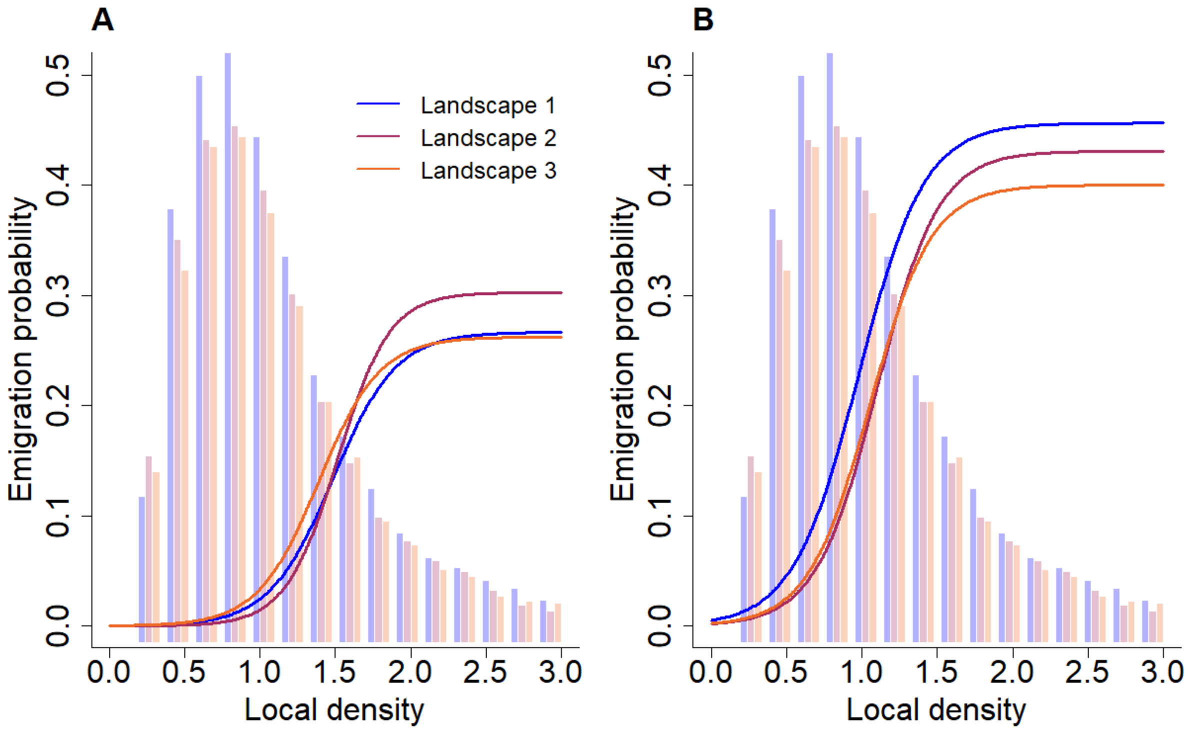

**Figure S1.** Emigration probability reaction norms (lines) for the average female (A) and male (B) after 2000 years of evolution on the three different landscapes. Bars represent frequencies of local densities from years 1800 to 2000.

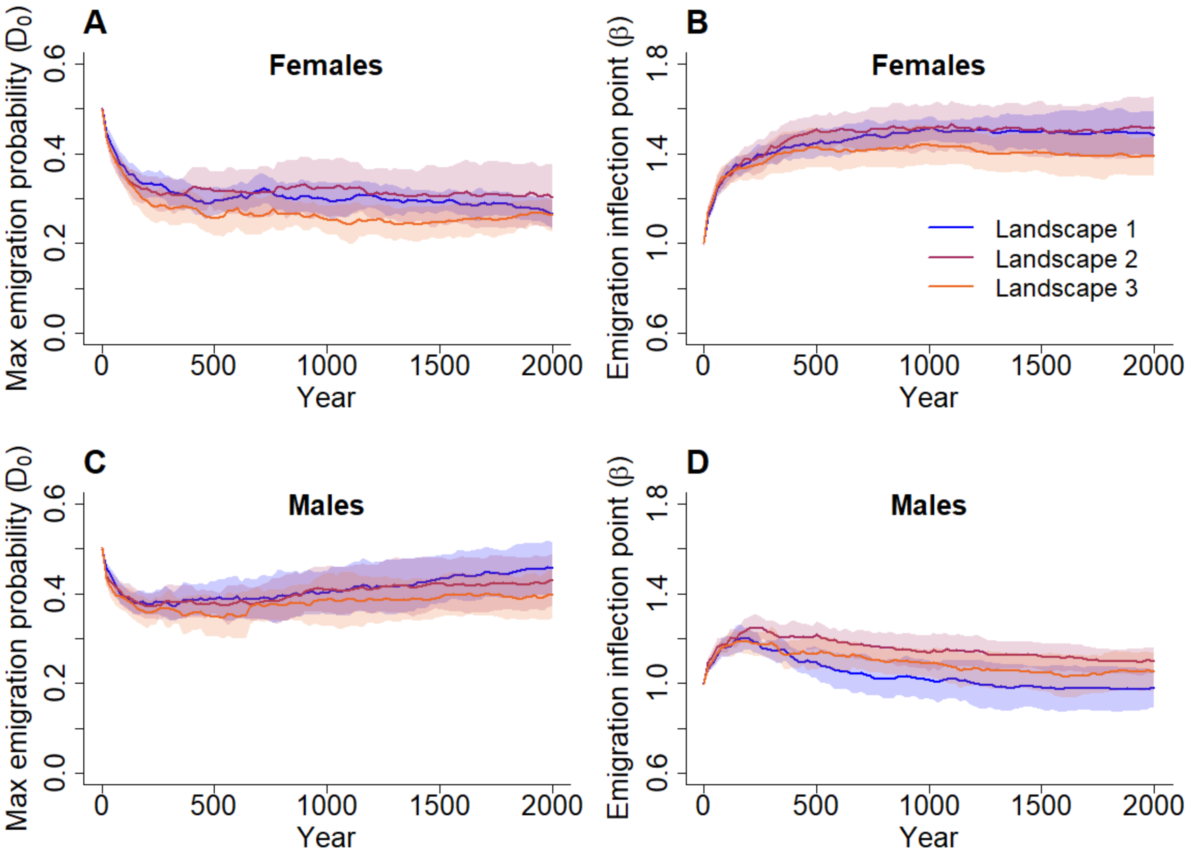

**Figure S2.** Evolution of mean female and male emigration traits in the three different landscapes. A) Mean female maximum emigration probability ($D_0$); B) mean female emigration inflection point ($\beta$); C) mean male $D_0$; D) mean male $\beta$. Mean trait values are averaged over 10 replicate simulations.

### ***Evolution of dispersal bias***

In RangeShifter 2.0, there is an option to include goal bias (GB), i.e. a tendency to move towards a particular destination (Aben et al., 2014), when modelling the transfer phase with SMS. However, as dispersers in RangeShifter are naïve and have no goal, it may be applied only in the 'negative' sense of moving away from the natal location, i.e. as a dispersal bias, which is implemented in a similar way to DP but relative to the direction from the natal site to the current location. The dispersal bias (DB) is further subject to a decay in strength as a function of the accumulated number of steps taken. This enables a dispersal path to follow a straighter trajectory initially, and later become more tortuous and responsive to perceived landscape costs. The dispersal bias applied for individual *i* at step *N* is determined by:

$$DB_{i,t} = 1 + \frac{GB_i - 1}{1 + e^{-(N - \beta_{DB,i})(-\alpha_{DB,i})}}$$

where $GB_i$, $\beta_{DB,i}$ and $\alpha_{DB,i}$ are the individual's phenotypes for the three evolving traits representing: the initial goal bias (GB), the inflection point ($\beta_{DB}$) and the slope at the inflection point ($\alpha_{DB}$).

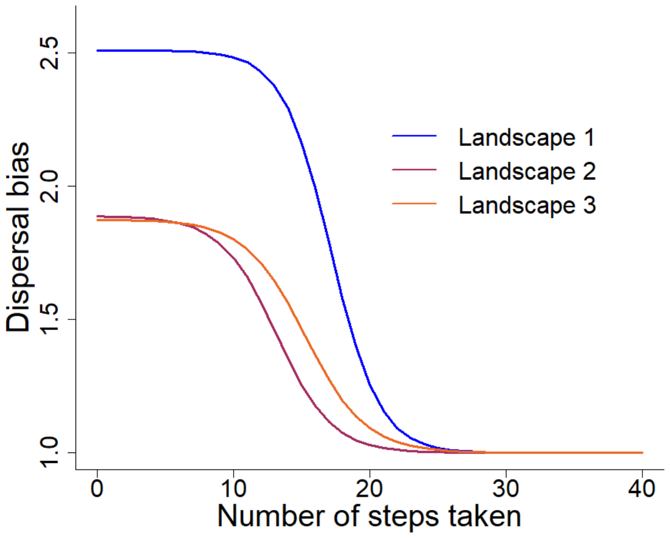

**Figure S3.** Dispersal bias decay curve for the average disperser after 2000 years of evolution.

***Evolution of density dependent settlement reaction norms***

The settlement probability for individual *i* in a patch it has reached is given by (see User Manual, section 2.5.5):

$$p_i = \frac{S_{0,i}}{1 + e^{-\left(\frac{N_{x,y}}{K_{x,y}} - \beta_{s,i}\right) * \alpha_{s,i}}}$$

where $S_{0,i}$, $\beta_{s,i}$ and $\alpha_{s,i}$ are the individual's phenotype for the three evolving traits representing: the maximum settlement probability ($S_0$), and the function inflection point ($\beta_s$) and slope at the inflection point ($\alpha_s$). $N_{x,y}$ and $K_{x,y}$ represent the patch total number of individuals and carrying capacity respectively. Males have the additional condition of requiring the presence of at least one female to settle in the patch.

The maximum settlement probability and the inflection point of the density-dependent settlement reaction norm evolved to relatively high mean values for both sexes in Landscape 1 (Fig. S5), so that almost all males and around 80% of females were likely to settle at the first suitable patch encountered unless it was substantially above carrying capacity (Fig. S4). When movement was inhibited (Landscapes 2 and 3), the inflection points for both sexes and the maximum probability for males evolved to slightly lower values (Fig. S4), and thus both sexes, but especially males, were somewhat less likely to settle in the first suitable patch encountered. The adoption of rather more selective settlement behaviour on more risky landscapes may appear counter-intuitive. However, the key here is that different dispersal traits do not evolve independently of each other. Disentangling the interactions between selection on these different dispersal rules requires much further work that this new version of RangeShifter can facilitate.

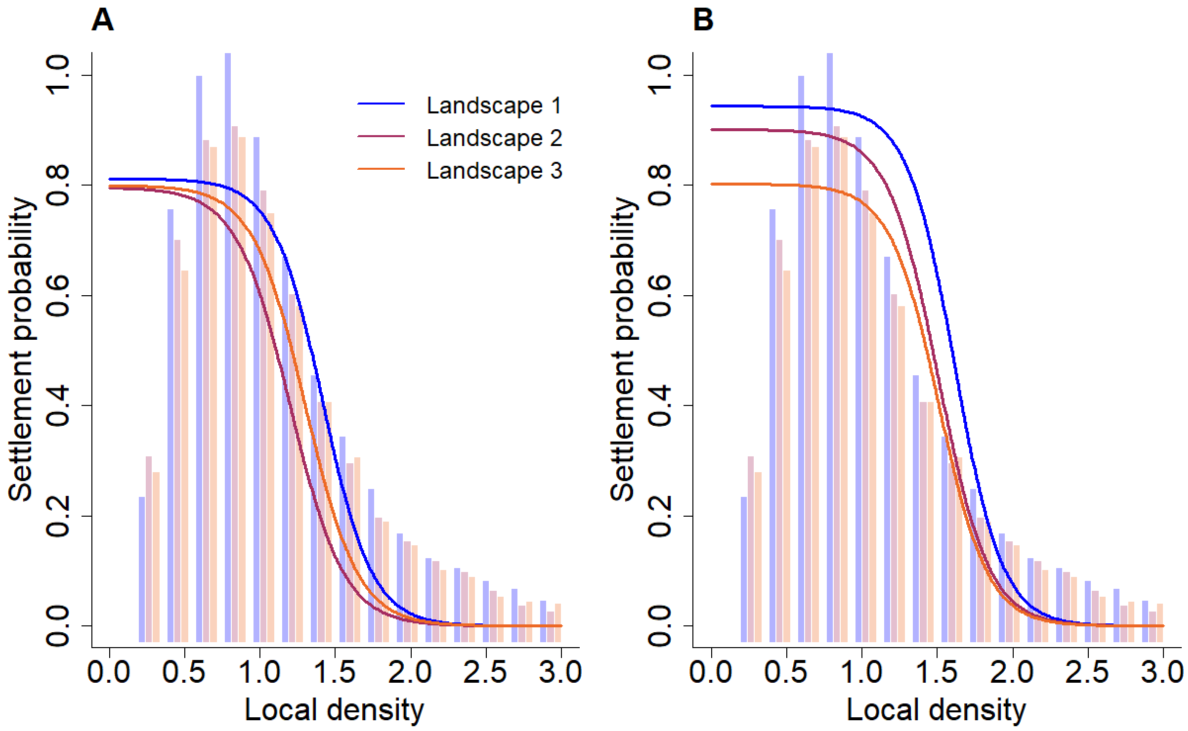

**Figure S4.** Settlement probability reaction norms (lines) for the average female (A) and male (B) after 2000 years of evolution on the three different landscapes. Bars represent frequencies of local densities from years 1800 to 2000.

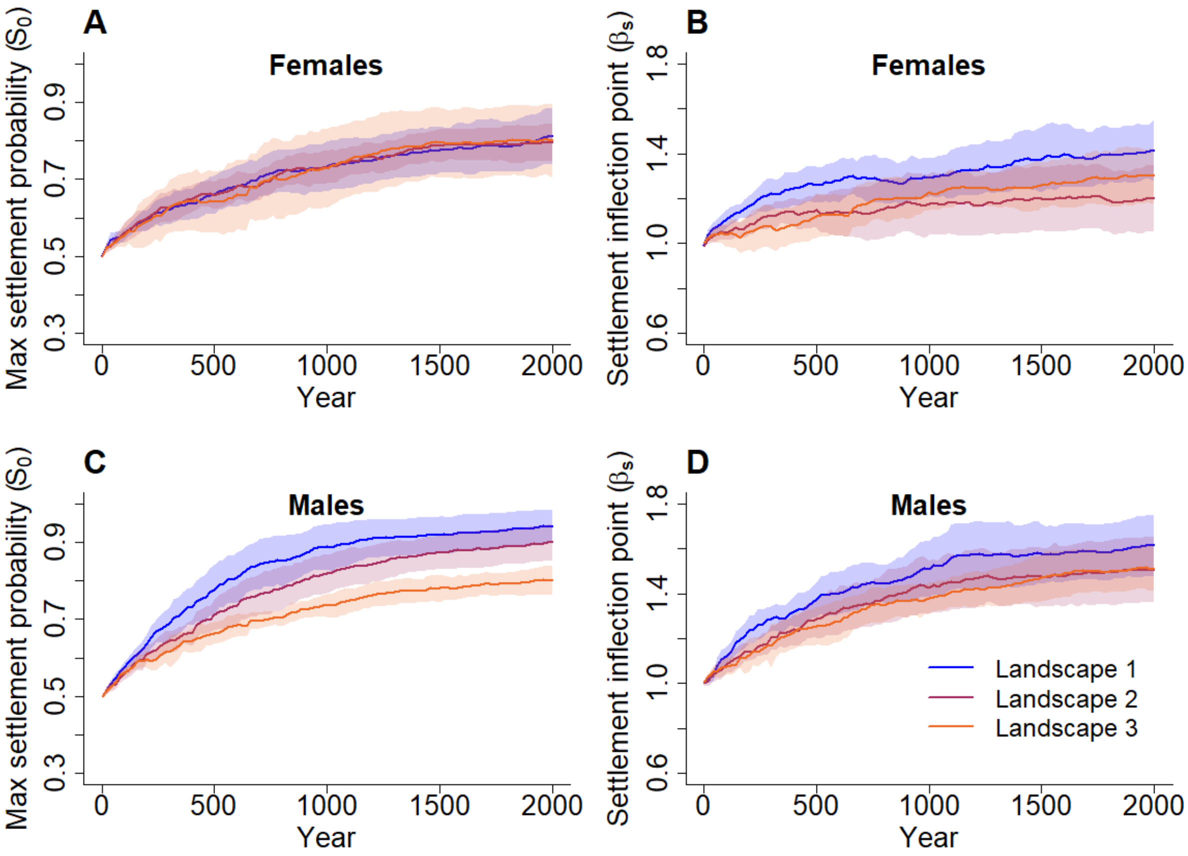

**Figure S5.** Evolution of mean female and male settlement traits in the three different landscapes. A) Mean female maximum settlement probability ($S_0$); B) mean female settlement inflection point ($\beta_s$); C) mean male $S_0$; D) mean male $\beta_s$. Mean trait values are averaged over 10 replicate simulations.

***Example dispersal paths***

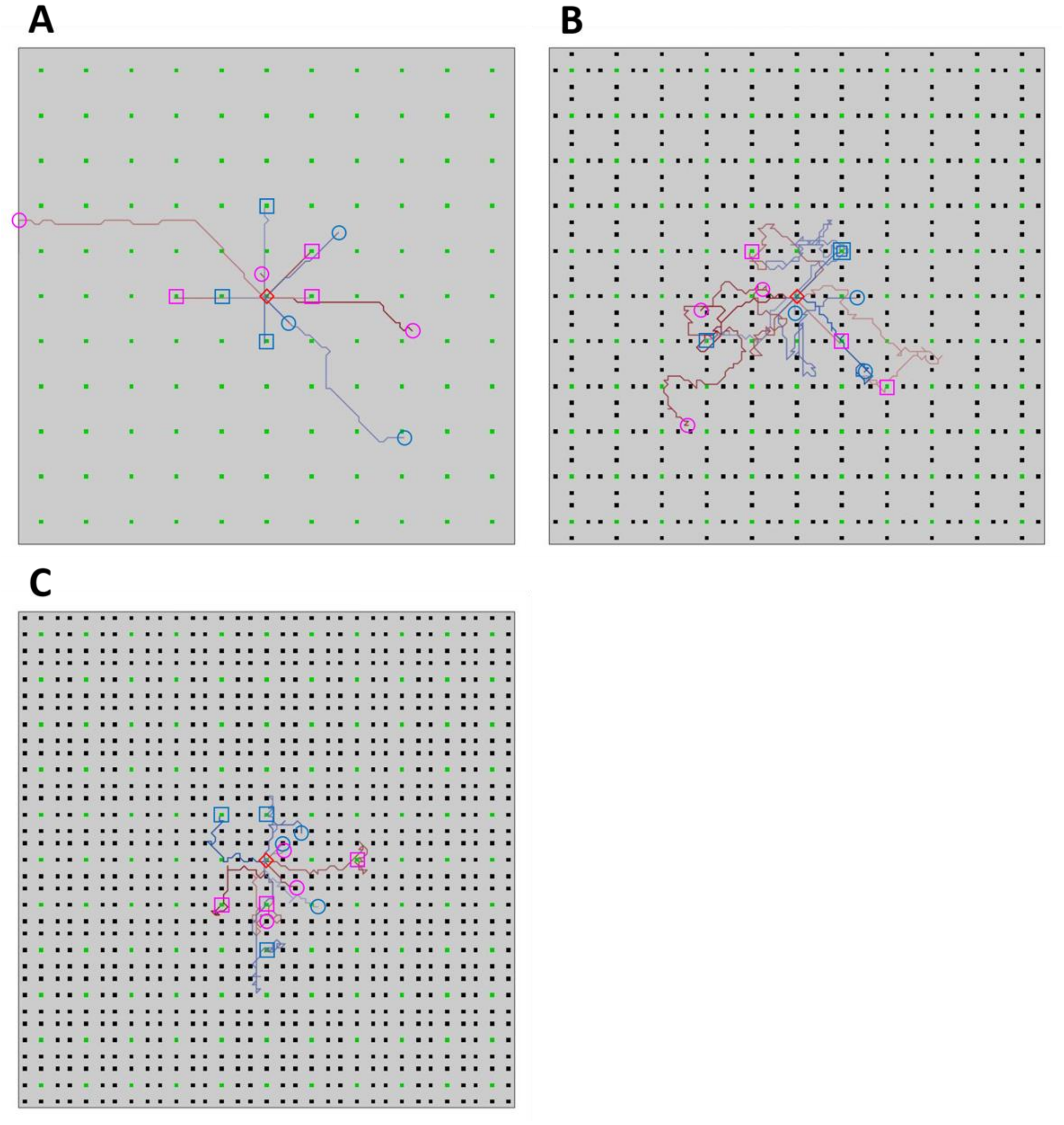

**Figure S6.** Example dispersal paths of individuals displaying the average SMS traits after 2000 years of evolution on landscapes having (a) no inhibitory cells (Landscape 1), (b) cells inhibitory to orthogonal movement (Landscape 2) and (c) cells inhibitory to orthogonal and diagonal movement (Landscape 3). Suitable cells are coloured green and inhibitory cells black. Six paths each of females (pink) and males (blue) are shown, all of which start at the central cell indicated by the red diamond. The end points of paths are indicated by squares if successfully reaching a suitable cell and by circles if the individual died in the matrix.
